# The Clp System in Malaria Parasites Degrades Essential Substrates to Regulate Plastid Biogenesis

**DOI:** 10.1101/718452

**Authors:** A. Florentin, D.R. Stephens, C.F. Brooks, R.P. Baptista, V Muralidharan

## Abstract

The human malaria parasite, *Plasmodium falciparum*, contains an essential plastid called the apicoplast. Most of apicoplast proteins are encoded by the nuclear genome and it is unclear how the plastid proteome is regulated. Here, we study an apicoplast-localized <u>c</u>aseinolytic-<u>p</u>rotease (Clp) system and how it regulates organelle proteostasis. Using null and conditional mutants, we demonstrated that the Clp protease (PfClpP) has robust enzymatic activity that is essential for apicoplast biogenesis. We developed a CRISPR/Cas9 based system to express catalytically-dead PfClpP, which showed that PfClpP oligomerizes as a zymogen and matured via trans-autocatalysis. The expression of a Clp chaperone (PfClpC) mutant led to the discovery of a functional chaperone-protease interaction essential for plastid function. Conditional mutants of the substrate-adaptor (PfClpS) demonstrated its essential function in plastid biogenesis. A combination of multiple affinity purification screens identified the Clp complex composition as well as putative Clp substrates. This comprehensive study reveals the molecular composition and interactions influencing the proteolytic function of the apicoplast Clp system and demonstrates its central role in the biogenesis of the plastid in malaria parasites.

## Introduction

The deadly malaria parasite, *Plasmodium falciparum*, has developed resistance to all currently available therapies, making the discovery of new drug targets a top priority^1^. Like most other apicomplexans, this eukaryotic parasite contains a plastid called the apicoplast^2^. This unique organelle evolved via a two-step endosymbiosis^3^; in the primary endosymbiotic event, a cyanobacterium was incorporated into a eukaryotic cell to form the modern chloroplast. During the secondary endosymbiotic event, a photosynthetic red alga was further taken up by a protist and led to the formation of a secondary plastid^4^. Although not photosynthetic, the apicoplast harbors essential prokaryotic metabolic pathways that are essential to the parasite throughout its complex life cycle^5^. In sharp contrast with its human host, the *Plasmodium* apicoplast shares molecular features with prokaryotes, plants and parasites, and therefore has the potential to encompass multiple parasite-specific drug targets^6,7^. Indeed, drugs that target apicoplast biology are in clinical use^8,9^. Most of those drugs (e.g. Doxycycline, Clindamycin) target the prokaryotic protein synthesis machinery in the apicoplast^10,11^. However, less than 10% of the apicoplast proteome is encoded by its own genome. The vast majority of the hundreds of apicoplast proteins are encoded in the nuclear genome and are transported to the organelle via the secretory pathway^12-14^. Available data suggest that the apicoplast does not control the translation of these nuclear encoded proteins, and, in fact, parasites without an apicoplast continue to express and accumulate these proteins in vesicle-like structures in the cytoplasm^15,16^.

Due to the inability of the apicoplast to control its own protein synthesis, it is likely that it maintains a stable proteome through protein degradation. This requires an organelle specific proteolytic machinery that has not yet been identified. We hypothesize that this function is executed by Clp (<u>C</u>aseino<u>l</u>ytic <u>p</u>rotease) proteins. This family of proteins consists of ClpP proteases that form multi-subunit proteolytic complexes, though the complex composition varies widely between different species and organelles^17,18^. The ClpP proteases associate with Clp ATPase chaperones that unfold and feed substrates into the ClpP barrel-like cavity for degradation^19,20^. In bacteria, they play pivotal roles in cell division, transport, stress response and virulence^21^. In plants chloroplasts, Clp proteins regulate the levels and activities of numerous metabolic enzymes and thus control chloroplast metabolism and differentiation^22^. Some of these metabolic pathways, such as isoprenoids biosynthesis, are conserved and essential in the apicoplast^4^.

Several putative Clp genes have been localized to the apicoplast of *P. falciparum*, but little is known about their roles in apicoplast biology or their essentiality for parasite asexual life cycle^23^. The putative *Plasmodium* Clp genes differ significantly from their bacterial orthologs and it is unclear whether they interact or even form a complex. They also include a putative noncatalytic subunit termed PfClpR that is absent in most bacteria^24^. We have previously shown that the *Plasmodium* ClpC chaperone (PfClpC) is essential for parasite viability and apicoplast biogenesis^16^. Here, we report the development and application of advanced molecular genetic tools in a clinically important non-model organism to gain detailed mechanistic insights into plastid Clp function and how this influences parasite biology. Collectively, these data demonstrate that the Clp complex and interactions within this proteolytic system function as an essential nexus regulating apicoplast biogenesis.

## Results and Discussion

### PfClpP is essential for apicoplast biogenesis and survival of malaria parasites

In order to assess the biological role of the *Plasmodium* ClpP homolog, PfClpP, and test its potential as a drug target, we attempted to generate a null mutant, replacing the *pfclpp* gene with a drug marker using CRISPR/Cas9 gene editing (Figure 1A). Despite PCR evidence for successful integration into the *pfclpp* locus (Supplementary Figure 1A), we repeatedly failed to retrieve live parasites following drug selection. This suggested that PfClpP is essential for parasite viability and we therefore decided to repeat the transfection in the presence of isopentenyl pyrophosphate (IPP). IPP is a small metabolite produced by the apicoplast which was shown to be the only essential function of the organelle during the asexual blood stages^15^. Transfection in the presence of IPP yielded live PfClpP knockout parasites (PfClpP^KO^) on the first attempt, and integration was confirmed by PCR (Figure 1B). Using an anti-PfClpP antibody we verified that PfClpP^KO^ mutants do not express PfClpP (Figure 1C). Immuno-fluorescence assays (IFA) revealed that apicoplast proteins such as Acyl Carrier Protein (ACP) and Chaperonin 60 (Cpn60) lost their typical apicoplast localization, and instead appear in vesicle-like structures (Figure 1D). These structures indicate damage to organelle integrity. Indeed, quantitative Real Time PCR (qRT-PCR) showed that the entire apicoplast genome disappeared in PfClpP^KO^ mutants, while the mitochondria genome was unaffected (Figure 1E). Consequently, removal of IPP from the culturing media resulted in the rapid death of the PfClpP^KO^ parasites (Figure 1F). Collectively, these data demonstrate that PfClpP is essential for parasite viability, and its function is required for the biogenesis of the apicoplast organelle.

**Figure 1.**
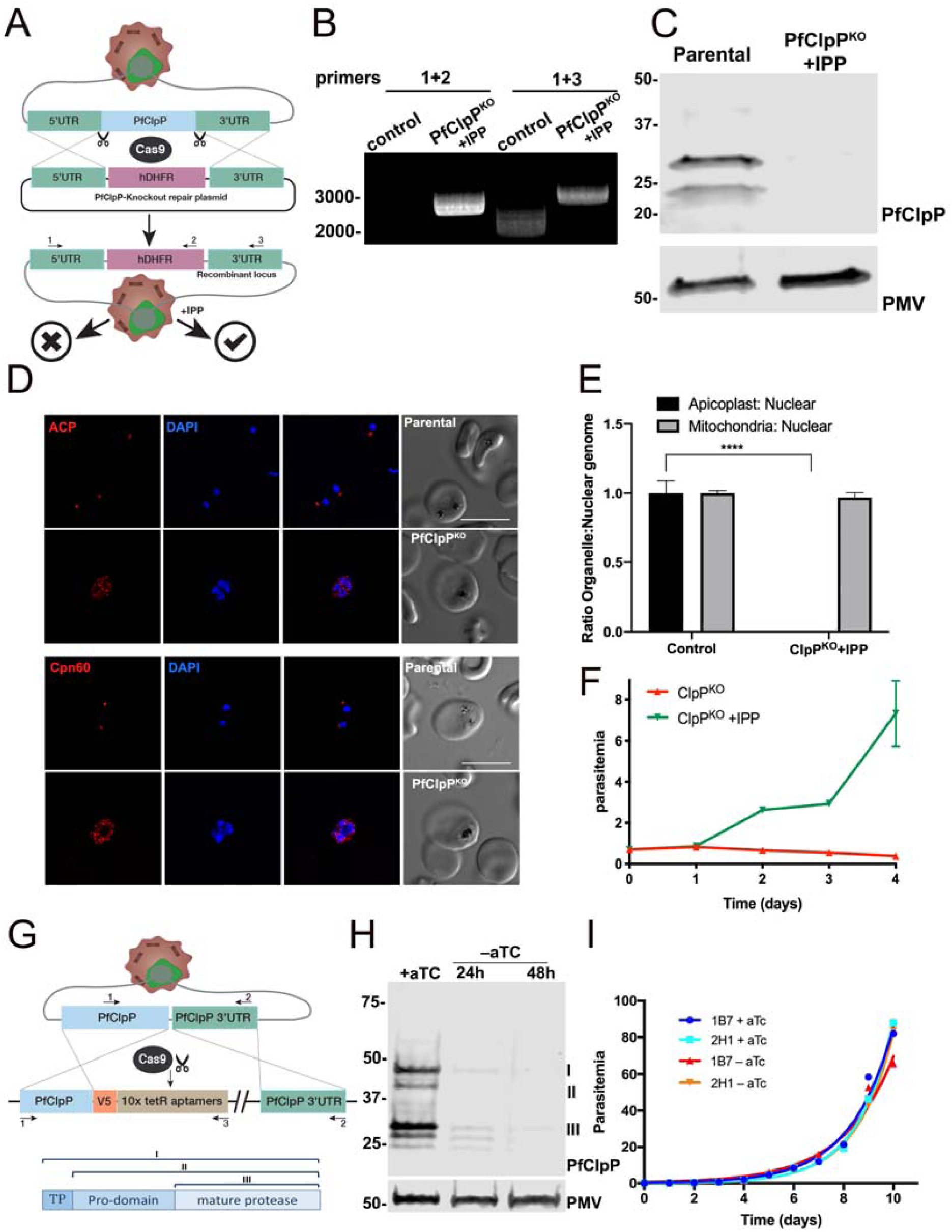
A. Generation of PfClpP knockout (PfClpP^KO^) parasites. CRISPR/Cas9 and two guide RNAs (gRNA) targeting the N- and the C-termini of PfClpP were used to facilitate integration. The repair plasmid contained 500bps homology regions to the 5’UTR and the 3’UTR of the *pfclpp* locus flanking the human dihydrofolate reductase selection marker (hDHFR). Through double crossover integration and drug selection, the gene was replaced with the drug marker. This strategy failed to retrieve live parasites unless IPP was added during drug selection. B. PCR test confirming hDHFR integration at the *pfclpp* locus in the presence of IPP. Genomic DNA was purified from transfected parasites. Primers 1+2 (see methods) were used to amplify an integration specific product. Primers 1+3 were used to amplify the region between the 5’UTR and 3’UTR of *pfclpp*, as illustrated in A. A shift of 1 KB corresponds to the integration of the hDHFR cassette. C. Western blot of parasite lysates from parental line and PfClpP^KO^ parasites probed with antibodies against PfClpP and Plasmepsin V (PMV, loading control), confirming loss of PfClpP expression in the null mutants. The protein marker sizes that co-migrated with the probed protein are shown on the left. D. Immunofluorescence imaging of fixed PfClpP^KO^ parasites stained with antibodies against ACP (Acyl Carrier Protein), and DAPI (upper panel) or Cpn60 (Chaperonin 60) and DAPI (lower panel). Both ACP and Cpn60 lose their typical apicoplast localization in PfClpP^KO^ and appear in vesicle-like structures. Z-stack images were deconvolved and projected as a combined single image. Representative images from biological replicates are shown. Scale bar, 5μm. E. Genomic DNA samples were collected from PfClpP^KO^ and control parasites for quantitative Real Time PCR analysis. Apicoplast: nuclear genome ratio was calculated for each sample. Mitochondria: nuclear genome ratio served as a control. Genome ratios were normalized to control parasites. Apicoplast genome is completely lost in PfClpP^KO^ mutants, while the mitochondria genome was unaffected. One representative experiment (out of 3 biological replicates) is shown. Data are represented as mean ± S.E.M (n=3 technical replicates, unpaired t-test, P value =0.00013) F. IPP was removed from PfClpP^KO^ parasites and parasitemia was monitored every 24 hours over 4 days via flow cytometry. Data are represented as mean ± S.E.M. (n=3 technical replicates). One representative experiment out of three biological replicates is shown. G. Diagram showing integration of the tetR-aptamer system into the *pfclpp* locus. The Cas9 nuclease together with a PfClpP-specific gRNA introduces a double-stranded break at the C-terminus of the *pfclpp* gene. The repair plasmid provides the template for double-crossover homologous recombination, introducing a 3xV5 tag and the tetR-Dozi cassette, including 10 aptamer repeats. H. PfClpP^apt^ parasites were incubated without aTc for 2 days, and lysates were isolated every 24 hr. Western blot shows parasite lysates probed with antibodies recognizing PfClpP (anti-V5) and Plasmepsin V (anti-PMV, loading control). The expected processing pattern of PfClpP is depicted: cytoplasmic fraction (I); apicoplast-localized zymogen (II) and mature protease after removal of the pro-domain (III). A significant reduction in PfClpP protein levels in achieved after 24 hours. The protein marker sizes that co-migrated with the probed protein are shown on the left. I. Two independent mutant clones (1B7 and 2H1) of PfClpP^apt^ parasites were grown with or without 0.5 μM aTc, and parasitemia was monitored every 24 hr over 11 days via flow cytometry. 100% of growth represents the highest value of calculated parasitemia (final parasitemia in the presence of aTc). Data are fit to an exponential growth equation and are represented as mean ± SEM (n= 3 technical replicates). One representative experiment out of three biological replicates is shown.

### PfClpP knockdown reveals robust enzymatic activity *in vivo*

Next, we wanted to investigate the molecular mechanisms of PfClpP function. To do that, we tagged the gene with a V5 tag and the *tetR*-aptamer conditional knockdown (KD) system, enabling translational repression of PfClpP expression when anhydrotetracycline (aTc) is removed from the culturing media^25^. Using CRISPR/Cas9, we incorporated the *tetR*-aptamer regulatory cassette at the 3’end of the *pfclpp* gene, creating the PfClpP^apt^ parasite line (Figure 1G). Using PCR analysis, we verified correct integration into the *pfclpp* locus (Supplementary Figure 1B), and by IFA we confirmed its apicoplast localization (Supplementary Figure 1C). Western blot analysis revealed the expected processing pattern of PfClpP; cytoplasmic fraction (I); apicoplast-localized zymogen (II) and a mature protease after proteolytic removal of the pro-domain (III) (Figure 1H)^16,23^. Inducing knockdown by removing aTc resulted in a significant reduction (>95%) in PfClpP protein levels, however this reduction had no effect on parasite growth (Figure 1H and 1I). This indicated that residual PfClpP protein levels were sufficient to maintain biologically functional enzymatic activity without any deleterious effects. The fact that PfClpP is essential for apicoplast function, and its maturation is proteolytically controlled, raised the possibility that it is post-translationally regulated.

### Development of a modular genetic system for *in vivo* expression

The study of such regulatory mechanisms *in vivo*, requires a genetic system that enables epistatic experiments in PfClpP^apt^ parasites. For this purpose, we developed a novel, modular tool for stable expression of tagged proteins in *P. falciparum*, that avoids the inconsistent overexpression that occurs when using episomal plasmids. Further, this expression system should be widely applicable in any parasite strain and can be deployed to study epistatic interactions in any parasite pathway. In this method, we use CRISPR/Cas9 editing to insert the gene of interest into a specific genomic locus, where it is expressed under an endogenous promoter. We chose the *pfhsp110c* locus, that encodes an essential cytoplasmic protein that is consistently expressed throughout the parasite life cycle^26^. To test whether the Hsp110 system can drive expression of apicoplast localized proteins, we introduced a 2A skip peptide at the end of the gene, followed by a GFP reporter with an apicoplast transit peptide (tp) derived from acyl carrier protein (ACP)^12^ (tp^ACP^-GFP, Figure 2A). Transfection and integration into the into the *pfhsp110c* locus were efficient, and using IFA we confirmed robust GFP expression in the apicoplast of all clones isolated (Figure 2B). Western blot analysis revealed the typical double-band, consisting of a weak upper band of the cytoplasmic fraction and a more prominent lower band of the apicoplast localized GFP after removal of the transit peptide (Figure 2C). We concluded that this method works well for expression of apicoplast proteins, and can be used to study genetic interactions between apicoplast Clp proteins.

**Figure 2.**
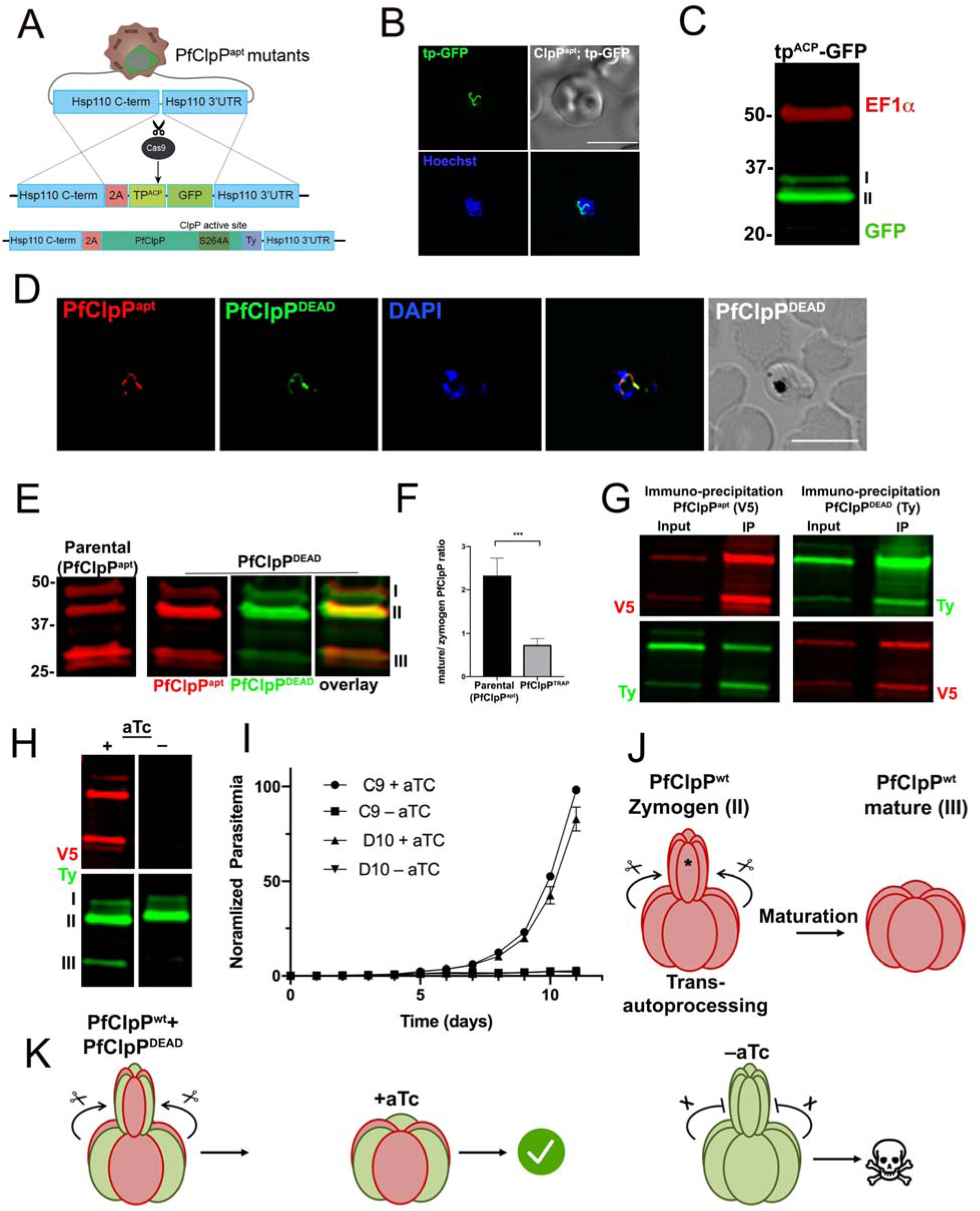
A. Diagram showing integration and expression through the Hsp110 expression system. Cas9 nuclease together with a gRNA targeting the *pfhsp110c* locus, introduces a double-stranded break at the c-terminus of the *pfHsp110c* gene. The repair plasmid provides two homology regions for homologous recombination, flanking a 2A skip peptide and a tagged gene of interest. Using this method, we introduced an apicoplast-localized GFP as a proof of concept, as well as a 3xTy-tagged catalytically-inactive PfClpP^DEAD^ variant with a point mutation in the protease active site (Ser264Ala). B. Live fluorescence microscopy visualizing the apicoplast localized GFP (tp^ACP^-GFP) expressed through the *pfHsp110c* locus in PfClpP^apt^ parasites. Parasite DNA was stained with Hoechst. Representative image from biological replicates is shown. C. Western blot of parasite lysates from tp^ACP^-GFP parasites probed with antibodies against GFP and EF1α (loading control). The protein marker sizes that co-migrated with the probed protein are shown on the left. D. Immunofluorescence microscopy of fixed PfClpP^apt^; PfClpP^DEAD^ parasites stained with antibodies against V5 (endogenous PfClpP^apt^, red), Ty (PfClpP^DEAD^, green) and DAPI (DNA). The two proteins co-localize in the apicoplast. Z-stack images were deconvolved and projected as a combined single image. Representative image from one of two biological replicates is shown. Scale bar, 5μm. E. Western blot of parental PfClpP^apt^ and PfClpP^apt^;PfClpP^DEAD^ parasite lysates probed with antibodies against V5 (endogenous *wild type* PfClpP, red) and Ty (PfClpP^DEAD^, green). The identical migration pattern indicates that PfClpP^apt^ and PfClpP^DEAD^ are similarly processed. The protein marker sizes that co-migrated with the probed protein are shown on the left. F. The ratio between the mature and zymogen forms of endogenous PfClpP (V5-tagged) is decreased upon PfClpP^DEAD^ expression, indicating that the dead variant interferes with the processing of the endogenous protease. Quantification of the ratio of II and III fractions of V5 tagged PfClpP from 3 biological replicates (p-value =0.0007, unpaired t-test). G. Co-Immunoprecipitation of PfClpP^apt^ (V5-tagged) and PfClpP^DEAD^ (Ty-tagged). 10^8^ parasites were isolated and pellets were incubated with either anti-V5 antibody-conjugated beads (left) or anti-Ty antibody-conjugated beads (right). Input and IP samples were loaded on SDS-page and blotted with anti-Ty and anti-V5 antibodies. The robust interaction between the two Clp protease variants suggests that they hetero-oligomerize into a mixed complex. H. PfClpP^apt^;PfClpP^DEAD^ parasites were incubated without aTc for 24 hours, and lysates were isolated. Western blot of these lysates was probed with antibodies against V5 (endogenous PfClpP, red) and Ty (PfClpP^DEAD^, green). PfClpP^apt^ knockdown results in the disappearance of the processed form of PfClpP^DEAD^, suggesting that the protease maturates through trans-autoprocessing. I. PfClpP^apt^;PfClpP^DEAD^ parasite clones (C9 and D10) were grown with or without aTc, and parasitemia was monitored every 24 hr over 11 days via flow cytometry. 100% of growth represents the highest value of calculated parasitemia (final parasitemia in the presence of aTc). Data are represented as mean ± SEM (n=3 technical replicates). One representative experiment out of two biological replicates is shown. J. A model illustrating the mechanisms of PfClpP oligomerization and maturation. PfClpP oligomerizes as a zymogen and subsequently matures by removing its pro-domain (denoted with *) via trans-autoprocessing. K. In the presence of the PfClpP^DEAD^ variant, a mixed complex is assembled and due to the robustness of PfClpP proteolytic activity, there is no effect on parasite growth. Upon aTc removal expression levels of endogenous PfClpP are dramatically reduced, and predominantly PfClpP^DEAD^ zymogen complexes are formed. Consequently, substrate degradation is inhibited and the parasites die.

### PfClpP oligomerizes as a zymogen and matures via trans-autoprocessing

The *Plasmodium* ClpP differs structurally from its bacterial orthologues primarily because it contains a transit peptide and a pro-domain (Figure 1H), both of which are removed at subsequent maturations steps. Like other proteins that are transported to the plastid, PfClpP possesses a transit peptide that is cleaved by a putative stromal peptide peptidase in the apicoplast^13^. A second cleavage event is required for the removal of the pro-domain through an unknown mechanism. It is possible that another protease is responsible for PfClpP pro-domain cleavage or that this is an autocatalytic event. Further, it is unknown how these processing events influence complex assembly and maturation of the apicoplast Clp protease, as well as other ClpP orthologues in well studied plastids, such as the chloroplast. To address all of these questions and to understand the mechanism of PfClpP oligomerization, we specifically interfered with the proteolytic activity of PfClpP *in vivo*.

For this purpose, we designed a PfClpP dead-protease mutant (termed PfClpP^DEAD^) with a point mutation in the protease active site (Ser264Ala) that renders it catalytically inactive^27^ (Figure 2A). It also includes the swapping of two residues (Glu308Arg/Arg285Glu) of an ion pair at the interface of the ClpP subunits^28^. The purpose of this switch is to reduce the affinity between PfClpP^DEAD^ and the endogenous PfClpP in order to minimize a potential dominant negative effect which may prevent us from studying PfClpP maturation and oligomerization. We used the Hsp110 expression system to express this Ty-tagged PfClpP^DEAD^ mutant in PfClpP^apt^ parasites (Figure 2A). The PfClpP^DEAD^ mutant co-localized in the apicoplast with PfClpP^apt^ (Figure 2D). Western blot analysis showed that the PfClpP^DEAD^ is processed similarly to the wild type PfClpP^apt^ to produce the cytoplasmic fraction (I); apicoplast-localized zymogen (II) and active protease without the pro-domain (III) (Figure 2E). However, a close examination revealed that the proteolytic maturation of the endogenous PfClpP^apt^ was reduced in the presence of the PfClpP^DEAD^ variant. In the parental PfClpP^apt^ parasite line, the majority of the apicoplast localized PfClpP zymogen was cleaved (fraction III, see ‘parental’ in Figure 2E). In contrast, the processing of the same endogenous V5 tagged PfClpP was inhibited in the PfClpP^DEAD^ parasites (Figure 2E). Quantification of the ratio between the processed (III) and the zymogen (II) fractions, revealed a decrease of about 70% in the proteolytic processing of PfClpP^apt^ (Figure 2F).

The shift in the processing rate could be explained by competitive binding between the two PfClpP alleles. To test this model, we co-immunoprecipitated the endogenous PfClpP^apt^ (V5 tag) and the PfClpP^DEAD^ (Ty tag). These experiments revealed that the two PfClpP alleles bind each other, indicating that they hetero-oligomerize into a mixed complex (Figure 2G). This was unexpected due to the swapping of the two residues (Glu308Arg/Arg285Glu) of the ion pair at the interface of the ClpP subunits. This cross-binding revealed several molecular features of PfClpP activation; the ability of each protease variant to co-immunoprecipitate not only the mature form (III) but also the zymogen (II) suggests that complex assembly precedes the final processing step of removing the pro-domain (Figure 2G and 2J).

### PfClpP protease activity is essential for complex function

The implication of the interaction between the wild-type and dead protease mutant became apparent during PfClpP^apt^ knockdown; reducing the levels of the endogenous active protease did not affect PfClpP^DEAD^ expression levels but it drastically inhibited its processing (Figure 2H). The inability of the parasites to remove the pro-domain from the dead protease in the absence of an active PfClpP, suggests that *in vivo* the PfClpP zymogen matures through trans-autoprocessing (Figure 2J). These data further suggest that the PfClpP zymogen (II) is inactive and this trans-autoprocessing is essential for PfClpP activation. These data support the model that PfClpP zymogen forms oligomers, which may be required for trans-autoprocessing and that the mechanism of maturation and oligomerization for plastid Clp proteases significantly differs from their bacterial counterparts. Whether these are general features of ClpP zymogens expressed in other plastids such as plant chloroplasts remains to be explored. Future structural studies with the ClpP dead zymogens may reveal additional features such as whether the zymogen is required to ensure proper trans-autocatalytic activity.

Importantly, the robust activity of PfClpP enabled the PfClpP^DEAD^ parasites to grow normally despite the presence of the mixed complexes. However, the presence of the PfClpP^DEAD^ mutant rendered them more sensitive to perturbations in PfClpP levels, and under these settings, knockdown of endogenous PfClpP^apt^ via removal of aTc resulted in parasite death (Figure 2I). Thus, the expression of PfClpP^DEAD^ in the presence of wild-type PfClpP revealed the mechanisms of PfClpP oligomerization and maturation (Figure 2J and 2K). The lethality of the dead-protease upon PfClpP knockdown showed that it is the capacity to degrade substrates that is required for organelle biogenesis and parasite survival (Figure 2K).

### PfClpC interacts with mature PfClpP in the apicoplast

Since bacterial ClpP orthologues are typically associated with Clp chaperones, we were interested to see whether the PfClpP complex interacts with any apicoplast chaperones. While the *Plasmodium* genome does not encode well-studied bacterial chaperones such as ClpA or ClpX, it does express an atypical AAA+ ATPase termed PfClpC with a putative tripeptide ClpP binding sequence^16,23^.

Using the Hsp110 expression system, we designed and expressed two isoforms of PfClpC in PfClpP^apt^ parasites (Figure 3A). The first was a copy of wild type PfClpC (PfClpC^wt^) with a C-terminal 3xTy tag. We transfected PfClpC^wt^, and confirmed its co-expression with PfClpP-V5 by western blot (Figure 3B, left lane) and their co-localization in the apicoplast by IFA (Figure 3E, upper panel). Importantly, PfClpP pulldown resulted in the co-immunoprecipitation of PfClpC^wt^, indicating a physical interaction between the chaperone and the protease (Figure 3C, left). The reciprocal PfClpC immunoprecipitation revealed that the chaperone interacts with the mature protease (III) but not with the zymogen form (II) or the full length PfClpP with the transit peptide (I) (Figure 3C, right). This indicates that PfClpC and PfClpP interact only upon co-localization in the apicoplast and that the chaperone binds the complex only after oligomerization and trans-autoproteolytic activation of the protease.

**Figure 3.**
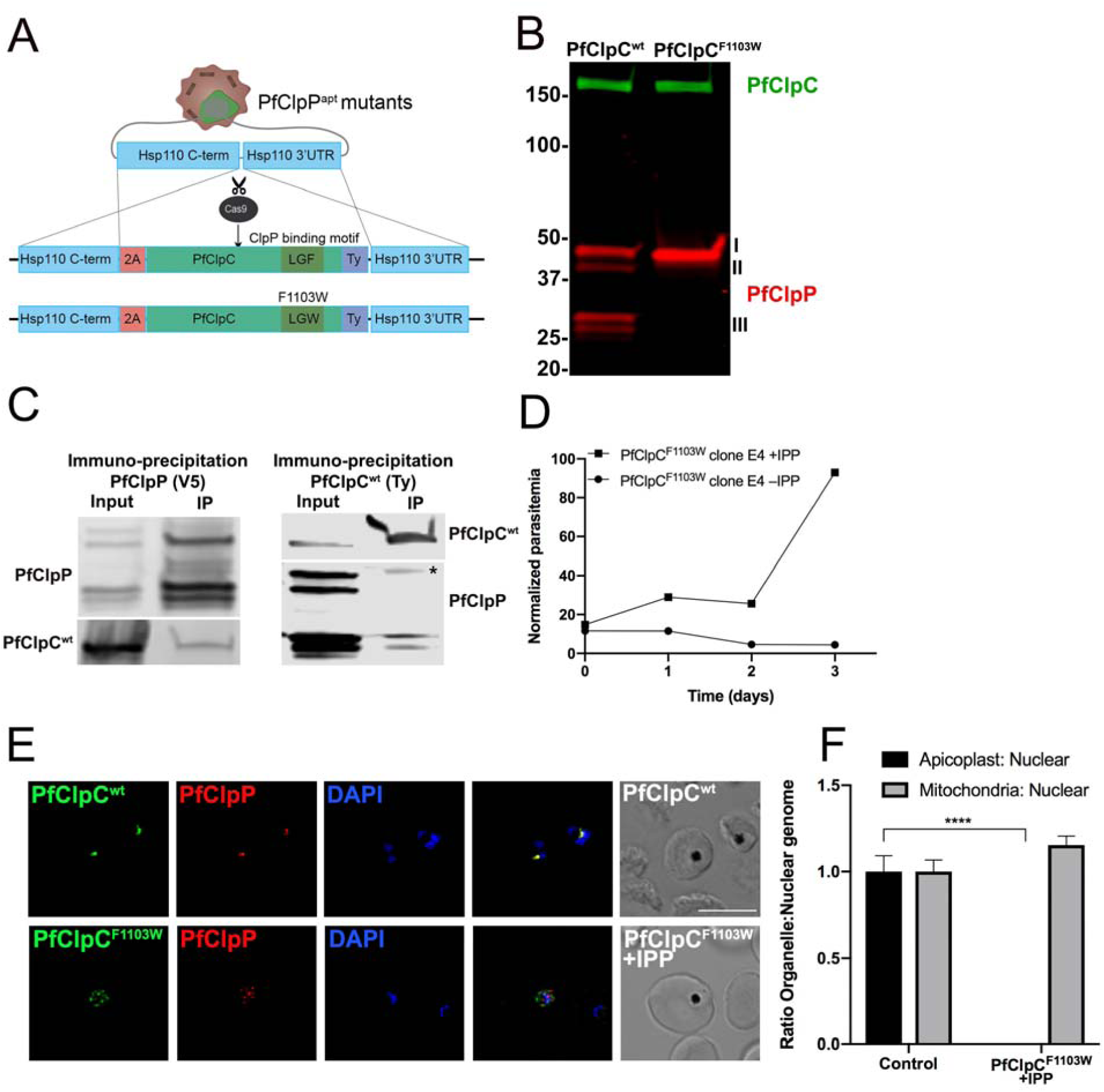
A. Diagram showing integration and expression of 3xTy-tagged PfClpP^DEAD^ variant into the *pfHsp110c* locus. Cas9 nuclease in complex with a gRNA targeting this locus introduces a double-stranded break at the C-terminus of the *pfHsp110c* gene. The repair plasmid provides two homology regions for homologous recombination, flanking a 2A skip peptide and a copy of wild type PfClpC tagged with 3xTy (PfClpC^wt^) or a PfClpC variant with a mutation in the putative PfClpP binding site (PfClpC^F1103W^). B. Western blot of parasite lysates from PfClpC^wt^ and PfClpC^F1103W^ parasites probed with antibodies against V5 (PfClpP, red) and Ty (PfClpC, green). Loss of all processing of PfClpP in PfClpC^F1103W^ parasites is due to apicoplast loss. The protein marker sizes that co-migrated with the probed protein are shown on the left. C. Co-Immunoprecipitation of PfClpP and PfClpC in PfClpP^apt^; PfClpC^wt^ parasites. 10^9^ parasites were isolated and pellets were incubated with either anti-V5 antibody-conjugated beads (PfClpP pulldown, left) or anti-Ty antibody-conjugated beads (PfClpC pulldown, rights). Input and IP samples were loaded on SDS-page and blotted with anti-Ty (PfClpC) and anti-V5 (PfClpP) antibodies. An interaction between the protease and the chaperone is observed. The residual heavy chain of the anti-Ty antibody that eluted with the IP sample despite the cross linker is also denoted (*). D. IPP was removed from PfClpC^F1103W^ parasites and parasitemia was monitored every 24 hours over 3 days via flow cytometry. 100% of growth represents the highest value of calculated parasitemia (final parasitemia in the presence of IPP). Data are represented as mean ± S.E.M. (n=3). One representative experiment out of two biological replicates is shown. E. Immunofluorescence microscopy of fixed PfClpC^wt^ and PfClpC^F1103W^ parasites stained with antibodies against V5 (PfClpP, red), Ty (PfClpC, green), and DAPI (DNA). The two proteins co-localize in the apicoplast in PfClpC^wt^ parasites but appear in vesicle-like structures in PfClpC^F1103W^ parasites. Z-stack images were deconvolved and projected as a combined single image. Representative images from one of two biological replicates are shown. Scale bar, 5μm. F. Genomic DNA samples were collected from PfClpC^F1103W^ and control parasites for quantitative Real Time PCR analysis. Apicoplast: nuclear genome ratio was calculated for each sample. Mitochondria: nuclear genome ratio served as a control. Genome ratios were normalized to control parasites. Data are represented as mean ± S.E.M (n=3 technical replicates, unpaired t-test, P value=0.00025). One representative experiment out of two biological replicates are shown.

### The Clp chaperone-protease interaction is essential for plastid biogenesis and parasite survival

The second PfClpC variant that we expressed via the Hsp110 system had a single point mutation in the 3^rd^ amino acid of a Leu-Gly-Phe motif (aa 1101-1103) that may be required for binding PfClpP (F1103W, Figure 3A)^20^. Aside from that mutation, this construct, termed PfClpC^F1103W^, was identical to the first (PfClpC^wt^) and was similarly tagged with 3xTy and expressed from the *pfhsp110c* locus in PfClpP^apt^ parasites, without interfering with the endogenous pfclpc gene (Figure 3A). Unexpectedly, multiple attempts to express the PfClpC^F1103W^ variant failed, suggesting that the mutated allele has a dominant negative effect on parasite viability. To rule out a general cytotoxic effect of transfection or any putative lethal consequences of interfering with PfHsp110c expression, we repeated transfections in the presence of the essential apicoplast metabolite, IPP. Transfection of PfClpC^F1103W^ in the presence of IPP has enabled us to retrieve live parasites. Western blot analysis revealed that in these parasites, all processing of PfClpP was abolished and the only observed band was the full-length cytoplasmic fraction (Figure 3B, right lane). Clones were isolated but their growth was entirely dependent on IPP, and removing it led to parasites death (Figure 3D). Using microscopy, we observed that both PfClpP^apt^ and PfClpC^F1103W^ did not appear in their typical apicoplast localization, and instead form vesicle-like structures indicative of apicoplast damage (Figure 3E, lower panel). Subsequently, quantitative real time PCR analysis revealed that the apicoplast genome disappeared from PfClpC^F1103W^ parasites (Figure 3F). These results indicate that interfering with the interaction between the PfClpC chaperone and the PfClpP protease inhibits complex function. Further, our data suggest that PfClpC dynamically interacts with PfClpP and this essential interaction may be required for degradation of substrates to ensure apicoplast biogenesis. To our knowledge, this is the first time that this interaction has been shown to be required for any plastid biogenesis.

### Tagging PfClpR with the tetR aptamer knockdown system confers a fitness cost

Another putative member of the plastid Clp complex and a potential regulator of Clp protease activity is ClpR, a non-catalytic subunit in the chloroplast Clp complex^23,24^. We therefore attempted to tag the apicoplast ortholog PfClpR with the V5 tag and the tetR aptamer knockdown system (Supplementary Figure 2A). Tagging *pfclpr* locus has been challenging^16^ and after multiple attempts, we finally succeeded in tagging the locus with the *tetR* aptamer. However, PCR analysis of the aptamer repeats region in PfClpR^apt^ parasites revealed that the number of aptamer repeats decreased during drug selection from 10 to 7 (Supplementary Figure 2B). This significantly compromised knockdown efficiency and prevented us from evaluating its effect on parasite replication (Supplementary Figure 2C). We concluded that tagging this locus confers a fitness cost and that, like PfClpP and PfClpC, PfClpR may be essential for plastid biogenesis. Nevertheless, the PfClpR gene was correctly tagged with the V5 epitope and, like PfClpP, was localized to the apicoplast (Supplementary Figure 2D). It is interesting to note that the non-catalytic PfClpR comprises an inactive Clp protease domain, but does not have a pro-domain, and migrates similarly on SDS-PAGE as the mature PfClpP (Supplementary Figure 2E).

### The adaptor protein PfClpS is essential for plastid biogenesis

Our data show that the proteolytic activity of the apicoplast Clp complex is central to its role in regulating plastid biogenesis. Therefore, it is essential to understand the mechanisms by which the apicoplast ClpP/R/C complex recognizes its substrates. Bioinformatic analysis have identified a putative substrate adaptor protein termed PfClpS, but its localization and physiological functions are yet to be determined^29^.

Therefore, we tagged PfClpS with a V5 tag and the *tetR*-aptamer knockdown system to generate PfClpS^apt^ parasites, and confirmed integration by PCR (Figure 4A). We observed co-localization of PfClpS and the apicoplast marker ACP (Figure 4D). Removal of aTc from PfClpS^apt^ parasites led to a significant protein knockdown (Figure 4B and 4C). As a consequence, aTc removal inhibited parasite growth, demonstrating that PfClpS is essential for parasite asexual replication (Figure 4E). Importantly, this growth inhibition was completely rescued by addition of IPP, linking the essential function of PfClpS to the apicoplast (Figure 4E). Microscopy revealed that apicoplast proteins such as ACP accumulated in vesicle-like structures, and quantitative Real Time PCR confirmed that the apicoplast genome disappears upon PfClpS knockdown (Figure 4D and 4F). Collectively, these data show that PfClpS activity is essential for parasite viability and, similar to other Clp proteins, it is required for apicoplast biogenesis.

**Figure 4.**
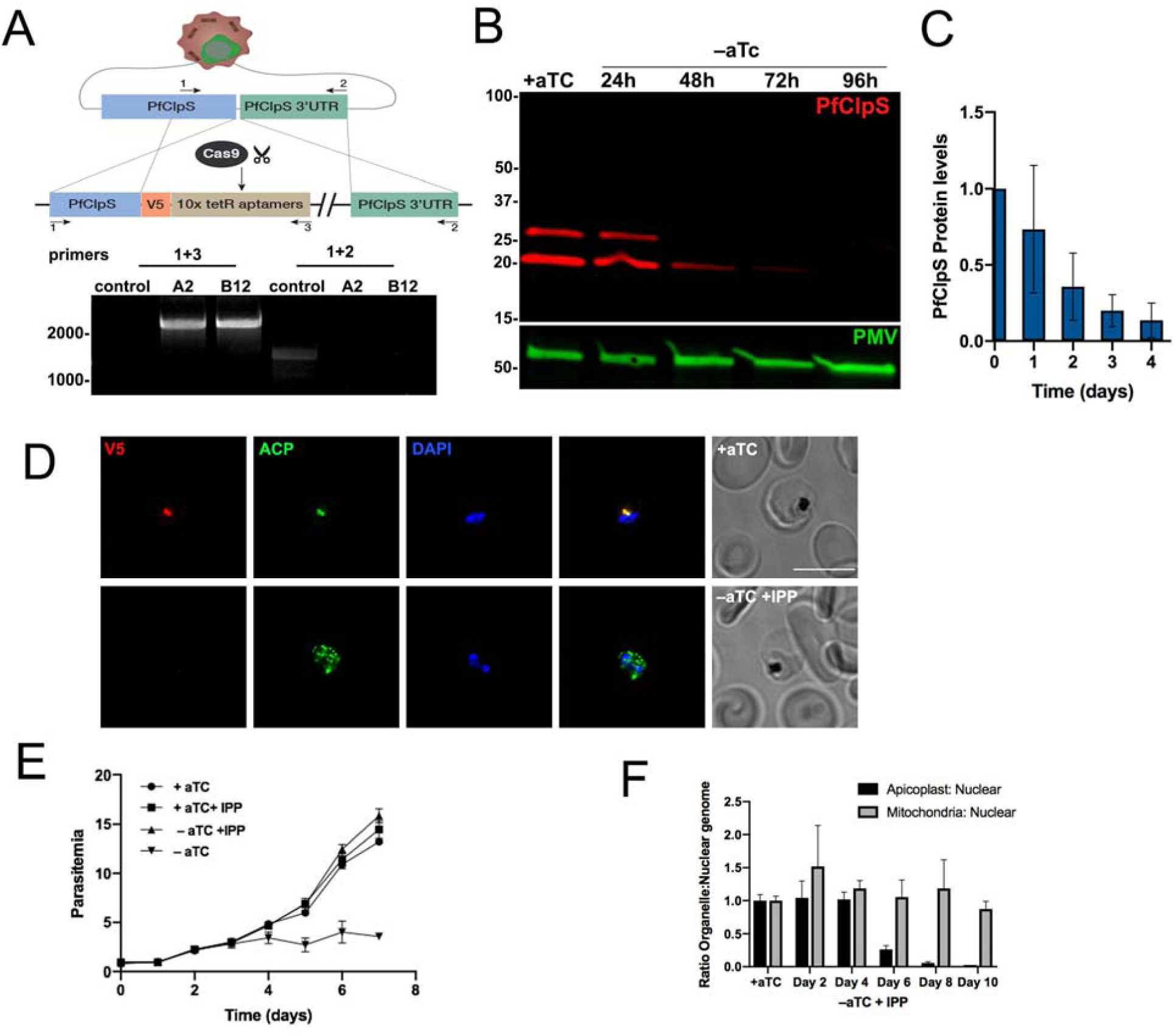
A. Upper panel: Diagram showing integration of the *tetR*-aptamer cassette into the *pfclps* locus. Cas9 nuclease with the gRNA introduces a double-stranded break at the beginning of the C-terminus of the *pfclps* gene. The repair plasmid provides homology regions for double-crossover homologous recombination, introducing a 3xV5 tag and the *tetR*-aptamer cassette. Lower panel: PCR test confirming *tetR*-aptamer integration at the *pfclps* locus. gDNA was purified from transfected parasites. Primers 1+2 or 1+3 (see methods) were used to amplify the native locus or an integration specific band, as illustrated in the diagram. B. PfClpS^apt^ parasites were incubated without aTc for 4 days, and lysates were isolated every 24 hr. Western blot of parasite lysates probed with antibodies against V5 (PfClpS, red) and Plasmepsin V (PMV, loading control, green). A significant reduction in PfClpS protein levels in apparent after 72 hours. The protein marker sizes that co-migrated with the probed protein are shown on the left. C. Quantification of PfClpS knockdown (as shown in B, n= 3 biological replicates). Values were normalized to PfClpS levels in the presence of aTc. D. Immunofluorescence microscopy of fixed PfClpS^apt^ parasites that were incubated for 8 days with aTc (upper panel) or without aTc and supplemented with IPP (lower panel). Parasites were stained with antibodies against V5 (PfClpS, red), ACP (Acyl Carrier Protein, green), and DAPI (DNA). PfClpS localizes to the apicoplast when aTc is present. Upon aTc removal, PfClpS disappears and ACP appears in vesicle-like structures. Z-stack images were deconvolved and projected as a combined single image. Representative images from one of two biological replicates are shown. Scale bar, 5μm. E. PfClpS^apt^ parasites (Clone A2) were grown with or without aTc and IPP, and parasitemia was monitored every 24 hr over 7 days via flow cytometry. Parasites grown without aTc exhibit a growth defect by day 4, and they die in the following days. Addition of IPP rescues the growth defect, indicating an apicoplast damage. 100% of growth represents the highest value of calculated parasitemia (final parasitemia in the presence of aTc). Data are represented as mean ± SEM (n= 3 technical replicates). One representative experiment out of three biological replicates is shown. F. Synchronized PfClpS^apt^ parasites were grown in the absence of aTc and presence of IPP. Genomic DNA samples were taken at the beginning of each replication cycle for real-time qPCR analysis. Apicoplast: nuclear genome ratio was calculated every 48h. Mitochondria: nuclear genome ratio served as a control. Genome ratios were normalized to parasites grown in the presence of aTc. Data are represented as mean ± SEM (n= 3 technical replicates). One representative experiment out of three biological replicates is shown.

### Reconstruction of the apicoplast Clp interactome

The phenotypic data that we collected so far, indicated a common biological function to the apicoplast Clp proteins. Moreover, the molecular mechanisms that we revealed here, including protease oligomerization and chaperone-protease interaction, further suggested the assembly of a functional proteolytic Clp complex. This has led us to test interactions between the other Clp proteins as well as to try and identify potential interactors, regulators and substrates.

For this aim, we performed immunoprecipitation (IP) of PfClpP^apt^ and PfClpP^DEAD^ parasites using V5 and Ty tags, respectively (supplementary Figure 3). For large scale proteomic analyses, potential Clp interactors were isolated from non-synchronized PfClpP^apt^ and PfClpP^DEAD^ parasites using anti-V5 and anti-Ty antibody-conjugated beads. The respective parental lines were used as controls, and all samples were analyzed by mass spectrometry. All detected proteins were filtered for predicted apicoplast localization and the abundance of each apicoplast-predicted protein was calculated and averaged between biological replicates. Using individual peptide abundance values, we calculated a ratio between protein abundance in IP and the parental control and set a threshold of ≥5-fold enrichment (Figure 5A, B). This resulted in a list of proteins that were either detected exclusively in the Clp IP or were at least 5-fold or more abundant than in the control.

**Figure 5.**
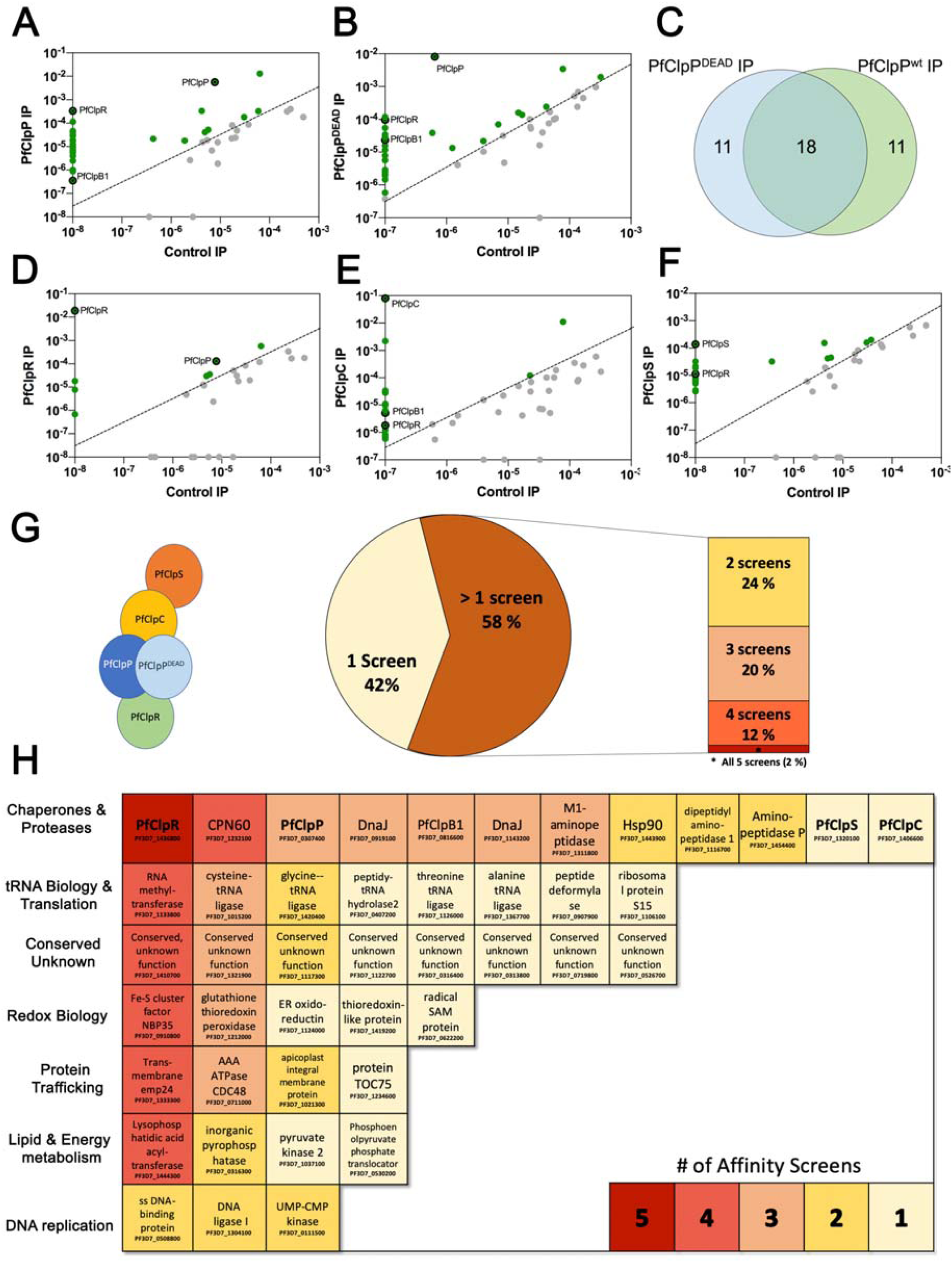
Mass spectrometry analysis of different Clp Immunoprecipitates isolated from the five distinct Clp-tagged parasite lines, using either V5 (for PfClpP^apt^/S^apt^/R^apt^) or Ty (for PfClpP^DEAD^/C^wt^) antibody-conjugated beads. All experiments were done using two biological replicates and the parental cell lines were used as a control (3D7 for PfClpP^apt^/S^apt^/R^apt^ and PfClpP^apt^ for PfClpP^DEAD^/C^wt^). All proteins detected in both biological replicates were filtered for predicted apicoplast localization. The abundance of each apicoplast-predicted protein was calculated and averaged between replicates. To filter for high confidence hits, we set a threshold of >5 fold enrichment in Clp IP vs. parental control. A. Mass spectrometry analysis of PfClpP Immunoprecipitate, isolated from PfClpP^apt^ parasite lysates. For PfClpP^apt^, 29 out of 45 proteins have passed the threshold and were defined high-confidence hits, including PfClpP, PfClpR and PfClpB1. B. Mass spectrometry analysis of PfClpP^DEAD^ Immunoprecipitate, isolated from PfClpP^apt^:PfClpP^DEAD^ parasite lysates. For PfClpP^DEAD^, 29 out of 49 proteins have passed the threshold and were defined high-confidence hits, including PfClpP, PfClpR and PfClpB1. C. Venn diagram showing the overlap between proteins identified in PfClpP^apt^ and PfClpP^DEAD^ pulldowns. The significant overlap increases the confidence in individual hits and suggests that the PfClpP^DEAD^ variant does not preferentially trap substrates. D. PfClpR was immunoprecipitated from lysates of PfClpR^apt^ parasites using anti V5 antibody-conjugated beads. Eight out of 29 PfClpR hits passed the threshold, and seven of them also appeared in the PfClpP^apt^ pulldown, including PfClpR and PfClpP. E. Mass spectrometry analysis of PfClpC^wt^ Immunoprecipitate, isolated from PfClpP^apt^:PfClpC^wt^ parasite lysates. For PfClpC^wt^, 18 out of 45 proteins have passed the threshold and were defined high-confidence hits, including PfClpC, PfClpR and PfClpB1. F. Mass spectrometry analysis of PfClpS immunoprecipitated from PfClpS^apt^ parasites. PfClpS IP yielded 30 apicoplast-localized proteins, out of which 20 have passed the threshold and were defined as high-confidence hits, including PfClpR and PfClpR. G. Analysis of all the hits that passed the thresholds in any of the 5 different screens generated a list of 50 apicoplast proteins that interact with at least one Clp member. A high degree of overlap reveals that most of the hits (58%) interact with at least two Clp proteins. H. The total Clp interactors from the five affinity screens were sorted into seven groups, based on predicted or reported biological functions. The groups include chaperone & proteases (including all Clp proteins), tRNA biology & translation, Redox biology, protein trafficking, lipids & energy metabolism and DNA replication and conserved proteins with unknown functions. PfClpR is the only protein that appeared in all five screens, but all groups (excluding DNA replication) contain proteins that appeared in four out of the five screens, demonstrating the role of the *Plasmodium* Clp complex as a master regulator of apicoplast biology.

Both types of PfClpP affinity screens (*wild type* and *DEAD* mutant) yielded similar numbers of proteins that are predicted to localize to the apicoplast (45 and 49, respectively). For both screens, 29 out of the total proteins passed the threshold and were considered high-confidence PfClpP interactors (Figure 5A,B and supplementary table 1). A significant overlap in the results of the two PfClpP screens, produced a list of 40 proteins that interact with either PfClpP variant (Figure 5C). This high degree of overlap in the hits suggested that the PfClpP^DEAD^ variant does not preferentially bind or trap substrates, which is to be expected due to the mixed nature of the heterocomplex (Figure 2G). Importantly, in both affinity screens we detected PfClpR as a high-confidence hit, as well as another Clp member, PfClpB1, which might be involved in protein refolding and quality control^30^.

In order to produce a comprehensive map of Clp interactome, we performed 3 more IP affinity screens, using PfClpR^apt^, PfClpS^apt^ and PfClpC^wt^ parasites lines. For PfClpR, out of 29 apicoplast-predicted proteins, only 8 passed the threshold and were considered high-confidence PfClpR interactors (Figure 5D and supplementary table 1). These eight interactors include PfClpP and PfClpR. Moreover, seven out of the eight PfClpR interactors were also appeared in the PfClpP screen, further suggesting that the pulldowns and bioinformatic analysis enrich for true Clp interactors.

Since our genetic and biochemical data indicate an interaction between the chaperone and the protease, we expected to find some overlap between PfClpC and the other screens. Similar to the protease pulldowns, we detected a total of 45 proteins in the PfClpC IP, and 18 of them passed the threshold and were considered high-confidence PfClpC interactors (Figure 5E and supplementary table 1). In addition to PfClpB1, another apicoplast chaperone that was shared between these screens was chaperonin 60 (Cpn60), which functions in folding of proteins transported to plastids^31^. The interaction of Cpn60 and PfClpB1 with the different Clp proteins suggests a protein quality control mechanism in the apicoplast to degrade damaged and misfolded proteins.

Lastly, to test the relationship between PfClpS and the Clp complex, as well as to enrich for potential Clp substrates, we used the PfClpS^apt^ parasites to perform the 5^th^ affinity screen. The immunoprecipitation and proteomic analyses were performed similarly to the other complex subunits. Out of 39 identified potential PfClpS interactors, 20 hits passed the high-confidence threshold, including PfClpR (Figure 5F and Supplementary Table 1). Due to the predicted function of PfClpS, we hypothesized that this list is enriched with putative Clp substrates. The PfClpS phenotypic data suggest that the substrates selected for degradation by PfClpS are essential for apicoplast biogenesis.

### Identification of apicoplast metabolic pathways that are regulated by the Clp system

Combining the high-confidence hits from the five affinity screens, resulted in a list of 50 potential Clp interactors (Figure 5G and Supplementary table 1). Interestingly, we found a high degree of overlap between the screens, with 58% of them shared between at least 2 different affinity screens (PfClpP/P^DEAD^R/C/S) (Figure 5G). This high level of overlap increased our confidence in the biological relevance of identified hits and allowed us to build an apicoplast Clp interactome. This sort of global map illuminates the composition of the complex as well as points to potential substrates and pathways that are regulated by the complex.

To do this, we sorted the total interactors from the five affinity screens into seven groups, based on predicted or reported biological functions (excluding six individual hits which, despite apicoplast transit-peptide prediction, were either experimentally-reported or functionally-predicted to localize to a different cellular compartment). The six biological functions were chaperone & proteases (including all Clp proteins), tRNA biology & translation, Redox biology, protein trafficking, lipids & energy metabolism and DNA replication (Figure 5H). The seventh group consisted of conserved proteins with unknown functions (Figure 5H). PfClpR was the only protein out of the 50 high-confidence hits that appeared in all five screens. Excluding DNA replication, each one of the biological groups contained proteins of high interest since they appeared in four out of the five screens (Figure 5H). Future work will determine whether specific interacting proteins function upstream (regulators) or downstream (substrates) of the Clp complex. Nevertheless, consistent with plant Clp systems, it is likely that many of the metabolic enzymes that were detected in the screens are Clp substrates that are regulated by the complex activity. The abundance of essential enzymes from very distinct metabolic pathways (DNA replication, RNA metabolism, lipid biosynthesis, etc.) demonstrates the role of the *Plasmodium* Clp complex as a master regulator of apicoplast biology.

### Proteostasis regulation by the apicoplast Clp complex

We propose a model in which the *Plasmodium* apicoplast maintains a stable proteome via protein degradation. This central function is required, among other things, for organelle biogenesis, and is executed and regulated by the proteolytic Clp complex (Figure 6A). As a consequence, interfering with the function of different subunits of the complex leads to organelle loss and parasite death. In the core of the Clp complex is PfClpP, a conserved potent serine protease (Figure 6B). PfClpP is transported to the apicoplast as a zymogen, which hetero-oligomerizes into a complex, and then matures into the active protease through trans-autocatalytic processing. PfClpR is a non-catalytic subunit that physically interacts with other complex subunits and may play a regulatory role. PfClpC is the chaperone that unfolds proteins and feed them into the proteolytic core. It interacts with the mature PfClpP but not with the zymogen, suggesting that it binds at a later step during complex assembly. Nevertheless, this interaction is crucial for complex activity and parasite survival. Finally, a small adaptor molecule, PfClpS, binds specific substrates and delivers them to the complex.

**Figure 6.**
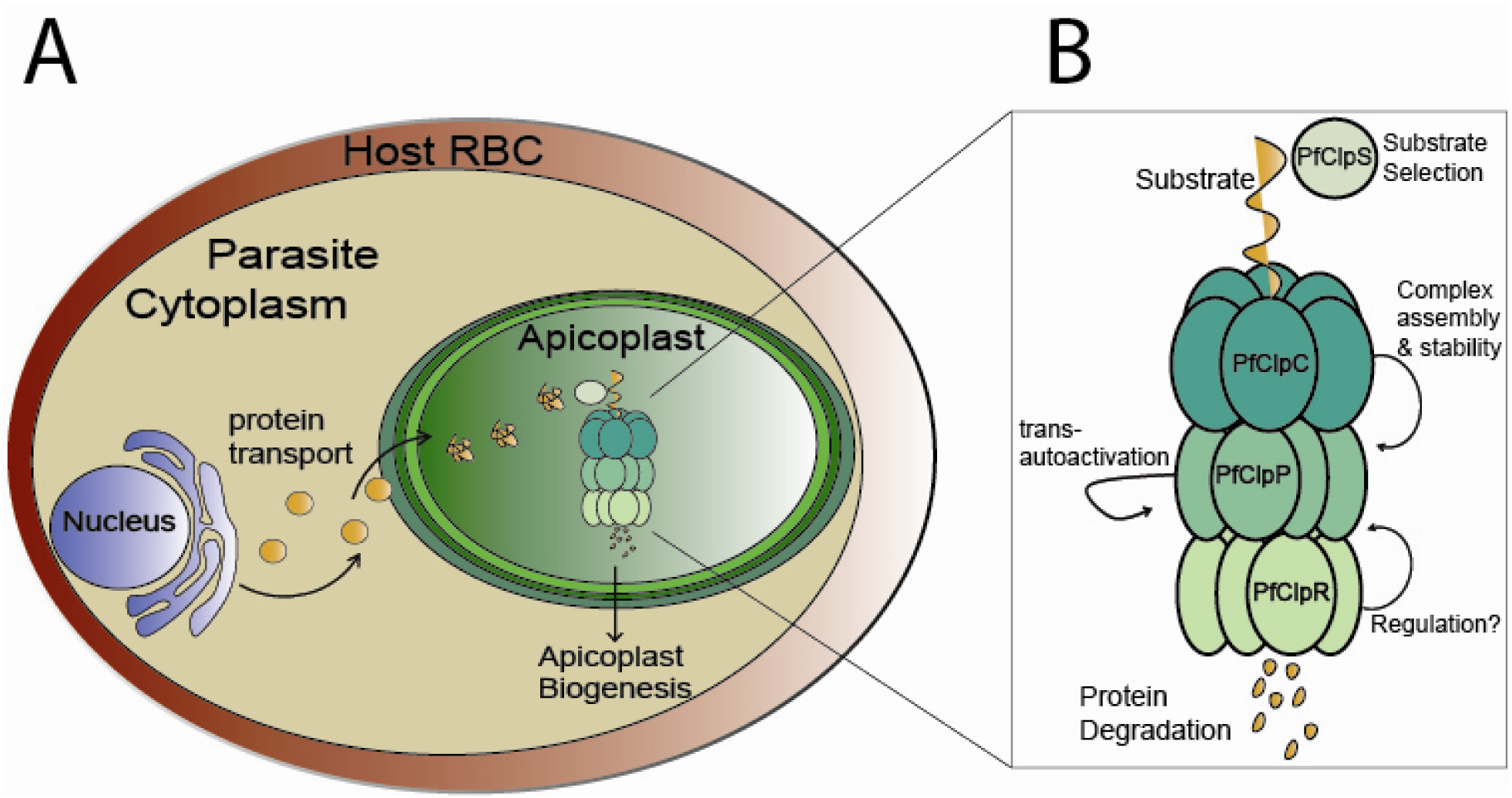
A. The Clp complex is a master regulator of apicoplast biology. Most of apicoplast proteins are encoded by the cell nucleus, and are transported to the organelle via the secretory pathway. According to this model, the apicoplast does not control their synthesis, and therefore a stable proteome in the organelle is via protein degradation. This essential function is required, among other things, for organelle biogenesis, and is executed and regulated by the proteolytic Clp complex. As a consequence, interfering with the function of different subunits of the complex leads to organelle loss and parasite death. B. Molecular function of the apicoplast Clp system. In the core of the Clp complex is PfClpP, a potent serine protease. PfClpP oligomerizes as a zymogen and then matures through trans-autoprocessing. PfClpR is a non-catalytic subunit that interacts with other Clp proteins and may have a regulatory role. PfClpC is the chaperone that unfolds proteins and feed them into the PfClpP proteolytic core. A transient interaction between PfClpC and the mature PfClpP is crucial for complex stability and function. A small adaptor molecule, PfClpS, binds and selects specific substrates for delivery to the complex.

Targeting protein synthesis in the apicoplast is clinically effective, as manifested by the clinical use of antibiotics such as doxycycline as antimalarials. However, such drugs inhibit the production of only a small fraction of apicoplast proteins, as the majority of them are transcribed by the cell nucleus and translated by the eukaryotic ribosomes in the parasite cytoplasm. Thus, a broader and potentially more efficient way to disrupt the apicoplast proteome would be to target its prokaryotic Clp degradation system. Indeed, due to their central function in prokaryotes, the bacterial ClpP homologs lie in the center of various drug discovery programs that identified potent inhibitors^32-36^. Ongoing work testing the effectiveness and specificity of these bacterial inhibitors against PfClpP may lead to the identification of antimalarials^16^. Our data and model suggest that small molecules inhibiting PfClpP activity will have a dual effect of reducing substrate cleavage as well as preventing the further production of the mature and active PfClpP. Future work is required to identify such potent inhibitors because our data show that, similar to its bacterial orthologs, the *Plasmodium* Clp complex is an excellent drug target.

## Materials &Methods

### Plasmids construction

Genomic DNA was isolated from *P. falciparum* using the QIAamp DNA blood kit (QIAGEN). All constructs utilized in this study were confirmed by sequencing. PCR products were inserted into the respective plasmids using Sequence and Ligation Independent Cloning (SLIC). Briefly, insert and cut vector were mixed with a T4 DNA polymerase and incubated for 2.5 minutes at room temperature, followed by 10 minutes incubation on ice and then transformed into bacteria. All restriction enzymes used in this study were purchased from New England Biolabs. All oligonucleotides used in this study are in Supplemental Table 2.

For the generation of PfClpP^KO^ donor plasmid, 500 bp from the 5’UTR of *pfclpp* (PF3D7_0307400) were amplified using primers 1+2 and 500 bp from the 3’UTR of *pfclpp* were amplified using primers 3+4. The human dihydrofolate reductase (hDHFR) drug selection cassette was amplified from pL6 vector using primers 5+6. All three pcr products were inserted into a TOPO cloning vector (ThermoFisher). Primer 7 was used in combination with primer 6 or 4 to test for accurate integration in the parasite following transfections. To increase efficiency of CRISPR/Cas9 mediated-integration, two guide RNAs were used, targeting both ends of the PfClpP gene. For expression of a C-term PfClpP guide RNA, oligos 8+9 were annealed and inserted into pUF1-Cas9-guide, and for expression of the N-term PfClpP guide RNA, oligos 10+11 were independently inserted into pUF1-Cas9-guide as previously described^16,37^.

For the generation of the PfClpP^apt^ conditional mutant, 500 bps from the 3’UTR and from the C-term of *pfclpp* gene were amplified pClpP-glmS vector^16^ using primers 18+19 and 20+21, respectively. The two products were conjugated together using PCR-sewing and inserted as one piece into pMG74-tetR-Dozi vector^25^. For expression of a C-term ClpP guide RNA, oligos 8+9 were annealed and inserted into pUF1-Cas9-guide.

For the generation of the PfClpR^apt^ conditional mutant, 500 bps from the 3’UTR and from the C-term of *pfclpr* gene (PF3D7_1436800) were amplified from pClpR-glmS vector^16^ using primers 33+34 and 35+21, respectively. The two products were conjugated together using PCR-sewing and inserted as one piece into pMG74-tetR-Dozi vector^25^. Primer 36 was used in combination with primer 34 or 38 to test for accurate integration in the parasite following transfections. Evaluating aptamer repeats numbers was done using primers 37+38. For expression of a C-term PfClpR guide RNA, oligos 39+40 were annealed and inserted into pUF1-Cas9-guide.

For the generation of the PfClpS^apt^ conditional mutant, 500 bps from the 3’UTR and from the C-term of *pfclps* gene (PF3D7_1320100) were amplified from gDNA using primers 41+42 and 43+44, respectively. The two products were conjugated together using PCR-sewing and inserted as one piece into pMG74-tetR-Dozi vector^25^. Primer 45 was used in combination with primer 42 or 38 to test for accurate integration in the parasite following transfections. For expression of a C-term PfClpS guide RNA, oligos 46+47 were annealed and inserted into pUF1-Cas9-guide. Primers 24+25 and 25+26 were used to test for accurate integration in the parasite following transfections.

For the generation of the Hsp110 expression system, we designed a repair plasmid that contains homology sequences from the *pfhsp110c* gene (PF3D7_0708800). The repair sequences include the last 429 bp (not including the stop codon) from the *pfhsp110c* gene, followed by a 2A skip peptide, a modular tagging cassette, and the first 400 bps from *pfhsp110c* 3’UTR. The modular tagging cassette comprised of a multiple cloning site with several optional tags including mCherry, V5, HA and Ty tags, allowing different tagging strategies using the same vector. The entire construct was created using gene synthesis (GeneScript) and was cloned into a puc57 backbone to create the puc57-Hsp110 vector. For expression of a PfHsp110c guide RNA, oligos 22+23 were annealed and inserted into pUF1-Cas9-guide.

For the generation of tp^ACP^-GFP parasites, tp^ACP^-GFP was amplified from pCEN-tp^ACP^-GFP using primers 27+28, and was inserted into puc57-Hsp110 vector (cut with MfeI and SpeI) using SLIC. For the generation of PfClpC^wt^ parasites, PfClpC (PF3D7_1406600) was amplified from gDNA using primers 29+32. To introduce the point mutation required for the generation of PfClpC^F1103W^ parasites, PfClpC was amplified from gDNA in two pieces, using primers 29+30 and 31+32. The two pieces were PCR-sewed together and inserted into puc57-Hsp110 vector (cut with MfeI and NheI) using SLIC.

For the generation of PfClpP^DEAD^ parasites, we introduced the following point mutations into the Open Reading Frame of PfClpP: Ser264Ala, Glu308Arg and Arg285Glu. The gene was synthesized (GeneScrpit) and was inserted into puc57-Hsp110 vector (cut with MfeI and NheI) using SLIC.

### Cell culture and transfections

Parasites were cultured in RPMI medium supplemented with Albumax I (Gibco) and transfected as described earlier^16,38,39^.

To generate PfClpP^KO^ parasites, a mix of three plasmids was transfected into 3D7 parasites in the presence or absence of 200uM Isopentynyl Pyrophosphate (IPP, Isoprenoids LC). The plasmid mix contained the PfClpP^KO^ donor plasmid and the two pUF1-Cas9-PfClpP-guide (N-terminal and C-terminal guides). Drug pressure was applied 48 hours post transfection using 2.5 nM WR99210 (Sigma) to select for parasites expressing hDHFR.

To generate PfClpP^apt^, PfClpR^apt^ and PfClpS^apt^ parasites, a mix of two plasmids was transfected into 3D7 parasites in the presence of 0.5 μM anhydrotetracycline (aTc, Cayman Chemicals). The plasmid mix contained 50 μg of pUF1-Cas9-guide (expressing the relevant guide: PfClpP/R/S) and 50 μg of the pMG74 based donor plasmid (with the related homology regions). Drug pressure was applied 48 hours post transfection using 2.5 μg/ml Blasticidin (BSD, Sigma). For each transfection, at least 2 clones were isolated via limiting dilution and used in all subsequent experiments.

To generate tp^ACP^-GFP, PfClpP^DEAD^, PfClpC^wt^ and PfClpC^F1103W^ parasites, a mix of two plasmids was transfected into PfClpP^apt^ parasites. For each different transfection the primers mix contained 2 plasmids; 50 μg of pUF1-Cas9-PfHsp110c-guide and 50 μg of the relevant marker-less puc57-Hsp110 repair plasmid (puc57-Hsp110-tp^ACP^-GFP/ puc57-Hsp110-PfClpP^DEAD^/ puc57-Hsp110-PfClpC^wt^/ puc57-Hsp110-PfClpC^F1103W^, respectively). For PfClpC^F1103W^, transfections were performed in the presence or absence of 200 μM IPP (Isoprenoids LC). Drug pressure was applied 48 hours post transfection, using 1μM DSM1 (BEI Resources), selecting only for Cas9 expression. DSM1 was removed from the culturing media once parasites clones were isolated using limiting dilution.

### Growth assays

For PfClpP^apt^, PfClpP^DEAD^, PfClpR^apt^ and PfClpS^apt^, asynchronous parasite cultures were washed 5 times and incubated without aTc. For PfClpP^KO^ and PfClpC^F1103W^, parasite cultures were washed once and incubated without IPP. Throughout the course of the experiment parasites were sub-cultured to maintain the parasitemia between 1-5% and parasitemia was monitored every 24 hours via flow cytometry. Cumulative parasitemia at each time point was back calculated based on actual parasitemia multiplied by the relevant dilution factors. Parasitemia in the presence of aTc or IPP at the end of each experiment was set as the highest relative parasitemia and was used to normalize parasites growth. Data were analyzed using Prism (GraphPad Software, Inc.)

For IPP rescue (PfClpS^apt^, PfClpP^KO^ and PfClpC^F1103W^) media was supplemented with 200 μM of IPP (Isoprenoids LC) in PBS.

### Western blot

Western blots were performed as described previously^40^. Briefly, parasites were collected and host red blood cells were permeabilized selectively by treatment with ice-cold 0.04% saponin in PBS for⍰10 min, followed by a wash in ice-cold PBS. Cells were lysed using RIPA buffer, sonicated, and cleared by centrifugation at 4°C. The antibodies used in this study were mouse anti-GFP, JL8 (Roche, 1:3000), mouse anti-Ty SAB4800032 (Sigma, 1:1000), rabbit anti-V5, D3H8Q (Cell Signaling, 1:1000), mouse anti-V5, TCM5 (eBioscience ™, 1:1000), mouse monoclonal anti-PMV (from D. Goldberg, 1:400), rabbit polyclonal anti-EF1α (from D. Goldberg, 1:2000) and rabbit polyclonal anti-PfClpP (from W. Houry, 1:4000). The secondary antibodies that were used are IRDye 680CW goat anti-rabbit IgG and IRDye 800CW goat anti-mouse IgG (LICOR Biosciences, 1:20,000). The Western blot images and quantifications were processed and analyzed using the Odyssey infrared imaging system software (LICOR Biosciences).

### Microscopy and image processing

For IFA cells were fixed using a mix of 4% Paraformaldehyde and 0.015% glutaraldehyde and permeabilized using 0.1% Triton-X100. Primary antibodies used are mouse anti Ty1 SAB4800032 (Sigma, 1:500), rabbit anti-V5, D3H8Q (Cell Signaling, 1:100), mouse anti-V5, TCM5 (eBioscience ™, 1:100), rabbit anti-Cpn60 (from B. Striepen, 1:1,000) and rabbit anti-ACP (from G. Mcfadden, 1:5,000). Secondary antibodies used are Alexa Fluor 488 and Alexa Fluor 546 (Life Technologies, 1:100). Cells were mounted on ProLong Diamond with DAPI (Invitrogen). For live cell imaging of tp^ACP^-GFP parasites were incubated with 8 μM Hoechst (ThermoFisher Scientific) in PBS. The imaging was performed using DeltaVision II microscope system with an Olympus IX-71 inverted microscope using a 100X objective. All images were collected as Z-stack, were deconvolved using the DVII acquisition software SoftWorx and displayed as maximum intensity projection. Image processing, analysis and display were preformed using SoftWorx and Adobe Photoshop. Adjustments to brightness and contrast were made for display purposes.

### Flow cytometry

Aliquots of parasite cultures (5μl) were stained with 8 μM Hoechst (ThermoFisher Scientific) in PBS. The fluorescence profiles of infected erythrocytes were measured by flow cytometry on a CytoFlex S (Beckman Coulter, Hialeah, Florida) and analyzed by FlowJo software (Treestar, Inc., Ashland, Oregon). The parasitemia data were fit to standard growth curve using Prism (GraphPad Software, Inc.).

### Quantitative Real Time PCR

Asynchronous parasites (PfClpP^KO^ and PfClpC^F1103W^) or Synchronized ring stage parasites (PfClpS^apt^) samples were collected and genomic DNA was purified using QIAamp blood kits (Qiagen). Primers that amplify segments from genes encoded by nuclear or organelles genomes were designed using RealTime qPCR Assay Entry (IDT). The following primer sequences were used: *cht1* (nuclear)-12+13; *tufA* (apicoplast)-14+15; *cytb3* (mitochondria)-16+17. Reactions contained template DNA, 0.5 μM of gene specific primers, and IQ ™ SYBR Green Supermix (BIORAD). Quantitative real-time PCR was carried out in triplicates and was performed at a 2-step reaction with 95°C denaturation and 56°C annealing and extension for 35 cycles on a CFX96 Real-Time System (BIORAD). Relative quantification of target genes was determined using Bio-Rad CFX manager 3.1 software. Standard curves for each primers set were obtained by using different dilutions of control gDNA as template, and were used to determine primers efficiency. The organelle: nuclear genome ratio of mutant parasites was calculated relative to that of the control. Unpaired t-test (Benjamini, Krieger, Yekutieli) was used to calculate significance between control and mutants.

### Immunoprecipitation

All pulldown experiments for proteomic analysis were performed in biological replicates using two different clones. For the pull-down proteomic analysis of PfClpP^apt^, PfClpR^apt^ and PfClpS^apt^ parasites, the parental line (wild type clone 3D7) was used a control and for PfClpP^DEAD^, PfClpC^wt^ the parental line PfClpP^apt^ was used a control. Immunoprecipitation protocols were performed using anti-V5 antibody (PfClpP^apt^, PfClpR^apt^ and PfClpS^apt^) or anti-Ty antibody (PfClpP^DEAD^ and PfClpC^wt^) as previously described^41^. Briefly, pellets from 10^9^ parasites were isolated using cold saponin and were lysed and sonicated in Extraction Buffer (40 mM Tris HCL pH 7.6, 150 mM KCL, and 1 mM EDTA) supplemented with 0.5% NP-40 (VWR) and HALT protease inhibitor (Thermo). 10% of the sample was kept for later analysis (input sample). Rabbit anti-V5 (Cell Signaling, D3H8Q) or anti-Ty (SAB4800032, Sigma) antibodies were crosslinked to Dynabeads protein G (Invitrogen) beads by incubating with 5 mM BS3 crosslinker (CovaChem) for 30 minutes. Quenching of crosslinker was performed using 1M Tris HCl (pH 7.5) for 30 minutes and then washing with 1.2M Glycine HCl (pH 2.5) to remove excess unbound antibody. Antibody-conjugated beads were then washed 3 time with PBS and then incubated with the supernatant at 4°C. Washes were performed using a magnetic rack (Life Technologies). Samples were run on SDS-page and gel slices were sent to mass spectrometry analyses.

For the co-Immunoprecipitation of PfClpP^DEAD^ and PfClpP^apt^, 2 samples of PfClpP^DEAD^ clone C9, each containing 10^8^ parasites were isolated as described above. Pellets were incubated with either anti-Ty antibody for PfClpP^DEAD^ (SAB4800032, Sigma) or with anti-V5 antibody for PfClpP^apt^ (Cell Signaling, D3H8Q). Input and IP samples were loaded on SDS-page and blotted with anti-Ty antibody (SAB4800032, Sigma) and with anti-V5 (Cell Signaling, D3H8Q). For the co-Immunoprecipitation of PfClpC^wt^ and PfClpP^apt^, 2 samples of PfClpC^wt^, each containing 10^9^ parasites were isolated as described above. Pellets were incubated with either anti-Ty antibody (SAB4800032, Sigma) or with anti-V5 (Cell Signaling, D3H8Q) antibody. Input and IP samples were loaded on SDS-page and blotted with anti-Ty antibody (SAB4800032, Sigma) or with anti-V5 (Cell Signaling, D3H8Q).

### Mass spectrometry and data analysis

PfClpP^apt^, PfClpR^apt^, PfClpS^apt^, PfClpP^DEAD^ and PfClpC^wt^ samples were sent to the proteomics shared resources in Fred Hutchinson Cancer Research Center and were run on the OrbiDEAD Elite. The data were searched using Proteome Discoverer 2.2 against the UP000001450 *Plasmodium falciparum* (Uniprot Nov 2018) database including common contaminants using Sequest HT and Percolator for validation. The search results were filtered for high confidence, with a strict 1% false discovery rate.

All detected proteins were analyzed for predicted apicoplast localization using four different bioinformatic algorithms: (i)PATS^42^; (ii)ApicoAP^43^; (iii)PlasmoAP^14^; and (iv) PlastNN^44^. We then compared results from all four algorithms and used known apicoplast proteins (such as Clp proteins) to estimate false-negative and false-positive rates for each algorithm. Based on these data, we chose to filter for apicoplast prediction using PlasmoAP, since it had a lower false-negative rate than PlastNN (higher sensitivity) and higher accuracy than the first two algorithms. The abundance of each apicoplast-predicted protein was calculated by summing the total intensities (MS1 values) of all matched peptides for each selected protein, and normalizing by the total summed intensity of all matched peptides in the sample, as previously described^44^. The resulted calculated value was then averaged between replicates. To filter for high confidence hits, we calculated the abundance ratio between Clp IP (PfClpP/ PfClpP^DEAD^/ PfClpR/ PfClpS/ PfClpC) and the parental control, and set a threshold of ≥5 fold enrichment.

## ACKNOWLEDGMENTS

We thank Belen Cassera for technical assistance, Geoffrey McFadden for anti-ACP antibody, Boris Striepen for anti-CPN60 antibody, Walid Houry for anti-PfClpP antibody, and Dan Goldberg for anti-PMV, and anti-EF1α antibodies; Julie Nelson at the CTEGD Cytometry Shared Resource Laboratory for help with flow cytometry and analysis; and Muthugapatti Kandasamy at the Biomedical Microscopy Core at the University of Georgia for help with microscopy. We acknowledge assistance of the Proteomics Resource at Fred Hutchinson Cancer Research Center for mass spectrometry. This work was supported the American Heart Association Postdoctoral Fellowship to A. F., and the U.S. National Institutes of Health (R21AI128195) to V. M.

---

**Supplementary Figure 1.**
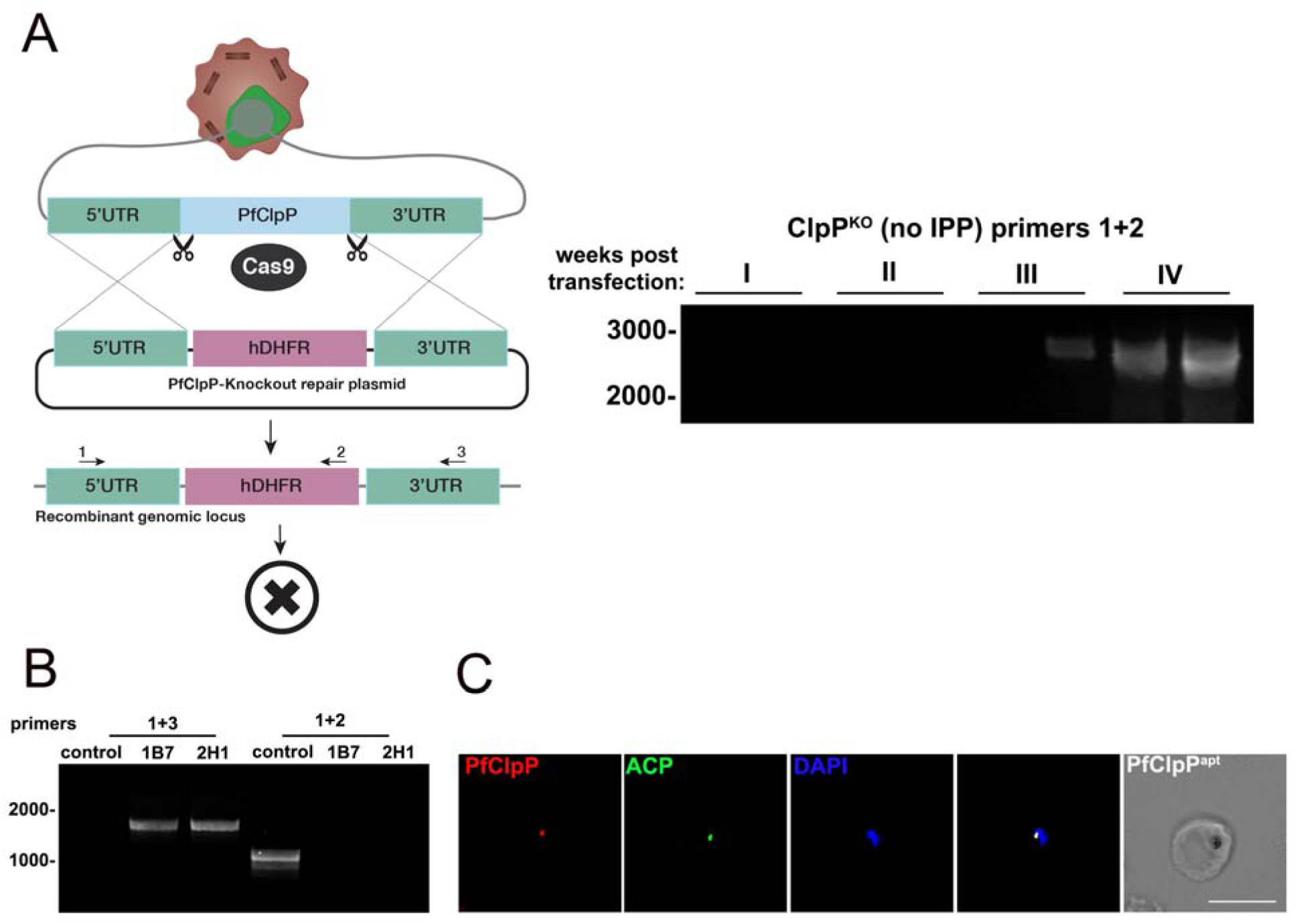
A. A diagram depicting the generation of PfClpP knockout (PfClpP^KO^) parasites. Cas9 nuclease and two guide RNAs targeting the N- and the C-termini of PfClpP were used to facilitate integration. The repair plasmid contained 500bps homology regions to the 5’UTR and the 3’UTR of the *pfclpp* locus flanking a drug marker (hDHFR, human Dihydrofolate Reductase). Through double crossover integration and drug selection, the gene was replaced with the drug marker. Despite evidence for integration into the genomic locus (bottom), this strategy failed to retrieve live parasites in the absence of IPP, indicating the PfClpP is essential to parasite viability due to its apicoplast function. B. PCR test confirming *tetR-Dozi* integration at the *pfclpp* locus. gDNA was purified from 2 independently selected mutant clones (1B7 and 2H1). Primers 1,2,3 (see methods) were used to amplify the native locus (1+2) or an integration specific product (1+3), as illustrated in the diagram in A. C. Immunofluorescence imaging of fixed PfClpP^apt^ parasites clone 1B7 stained with antibodies recognizing PfClpP (anti-V5, red), ACP (anti-Acyl Carrier Protein, green), and DAPI. Z-stack images were deconvolved and projected as a combined single image. Representative images from three biological replicates are shown. Scale bar, 5μm.

**Supplementary Figure 2.**
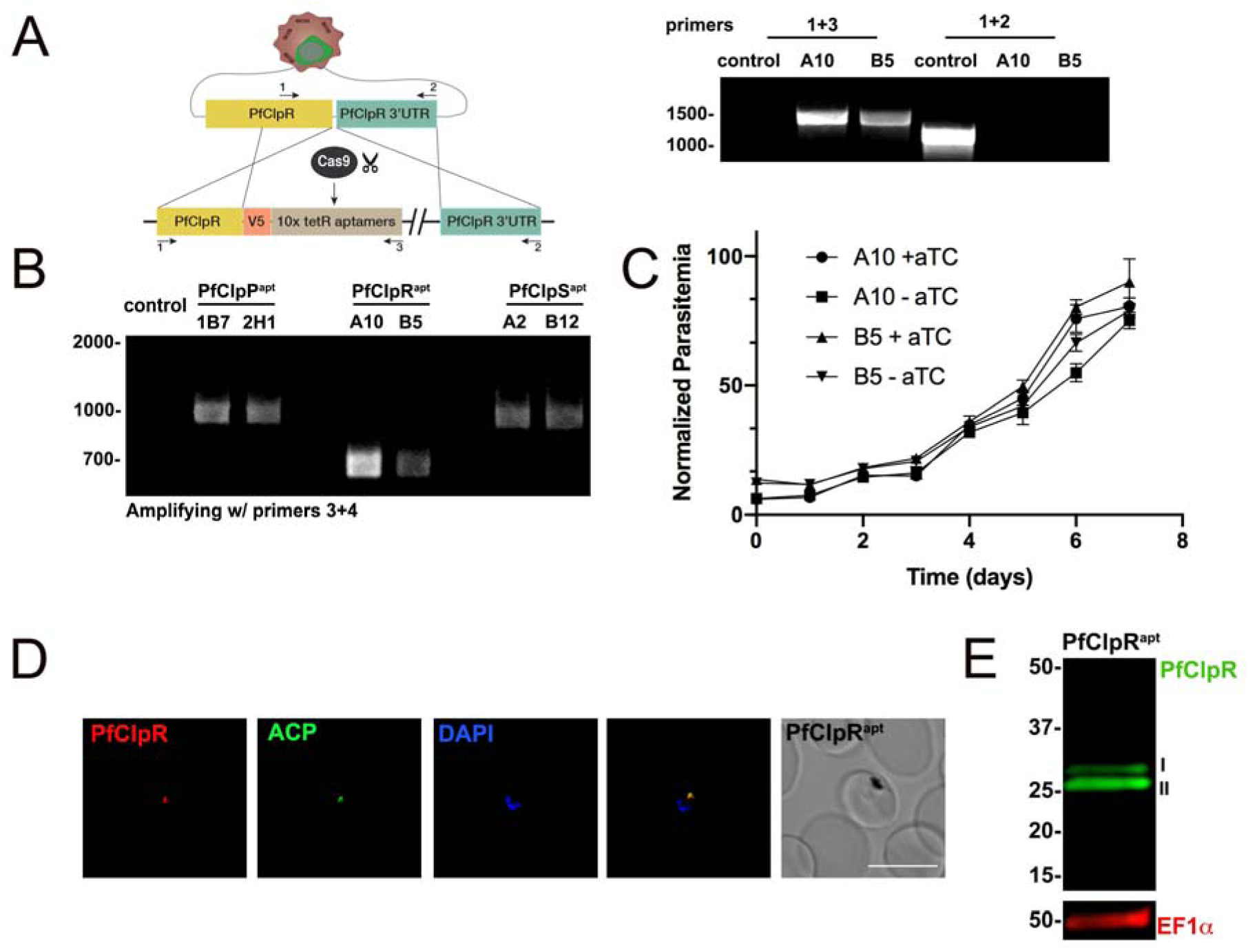
A. Diagram showing integration of the *tetR*-aptamer system into the *pfclpr* locus. Cas9 nuclease with a specific guide RNA introduces a double-stranded break at the C-terminus of the *pfclpr* gene. The repair plasmid provides homology regions for double-crossover homologous recombination, introducing a 3xV5 tag and the *tetR*-aptamer cassette. PCR test confirming *tetR*-aptamer integration at the *pfclpr* locus. Genomic DNA was purified from 2 mutant clones (A10 and B5). Primers 1,2,3 (see methods) were used to amplify the native locus (1+2) or an integration specific product (1+3), as illustrated in the diagram. B. PCR test to evaluate aptamer repeat numbers in *tetR*-aptamer cassette in genomic DNA isolated from PfClpP^apt^, PfClpR^apt^, and PfClpS^apt^ clones. The original plasmid contained 10 aptamer repeats, each 100pb long. Reduction in repeats numbers implies decreased binding by the tet-Repressor which compromises knockdown efficiency. While PfClpP^apt^ and PfClpS^apt^ clones show the expected 1 kb PCR product which corresponds to 10 aptamer repeats, all PfClpR^apt^ clones show a 0.7 kb product, indicating the loss of 3 aptamer repeats. C. Two PfClpR^apt^ clones (A5 and B10) were grown with or without 0.5 uM aTc, and parasitemia was monitored every 24 hr over 11 days via flow cytometry. 100% of growth represents the highest value of calculated parasitemia (final parasitemia in the presence of aTc). Data are represented as mean ± SEM (n= 3 technical replicates). One representative experiment out of two biological replicates is shown. D. Immunofluorescence microscopy of fixed PfClpR^apt^ parasites stained with antibodies against V5 (PfClpR^apt^, red), ACP (Acyl Carrier Protein, green), and DAPI (DAPI). Z-stack images were deconvolved and projected as a combined single image. Representative images from one of two biological replicates are shown. Scale bar, 5μm. E. PfClpR^apt^ lysates were isolated and probed with antibodies against V5 (PfClpR^apt^, red), and PMV (loading control). The doublet observed for PfClpR is typical for apicoplast proteins and include a cytoplasmic fraction (I) and an apicoplast-localized protein after the transit peptide had been removed (II). The protein marker sizes that co-migrated with the probed protein are shown on the left.

**Supplementary Figure 3.**
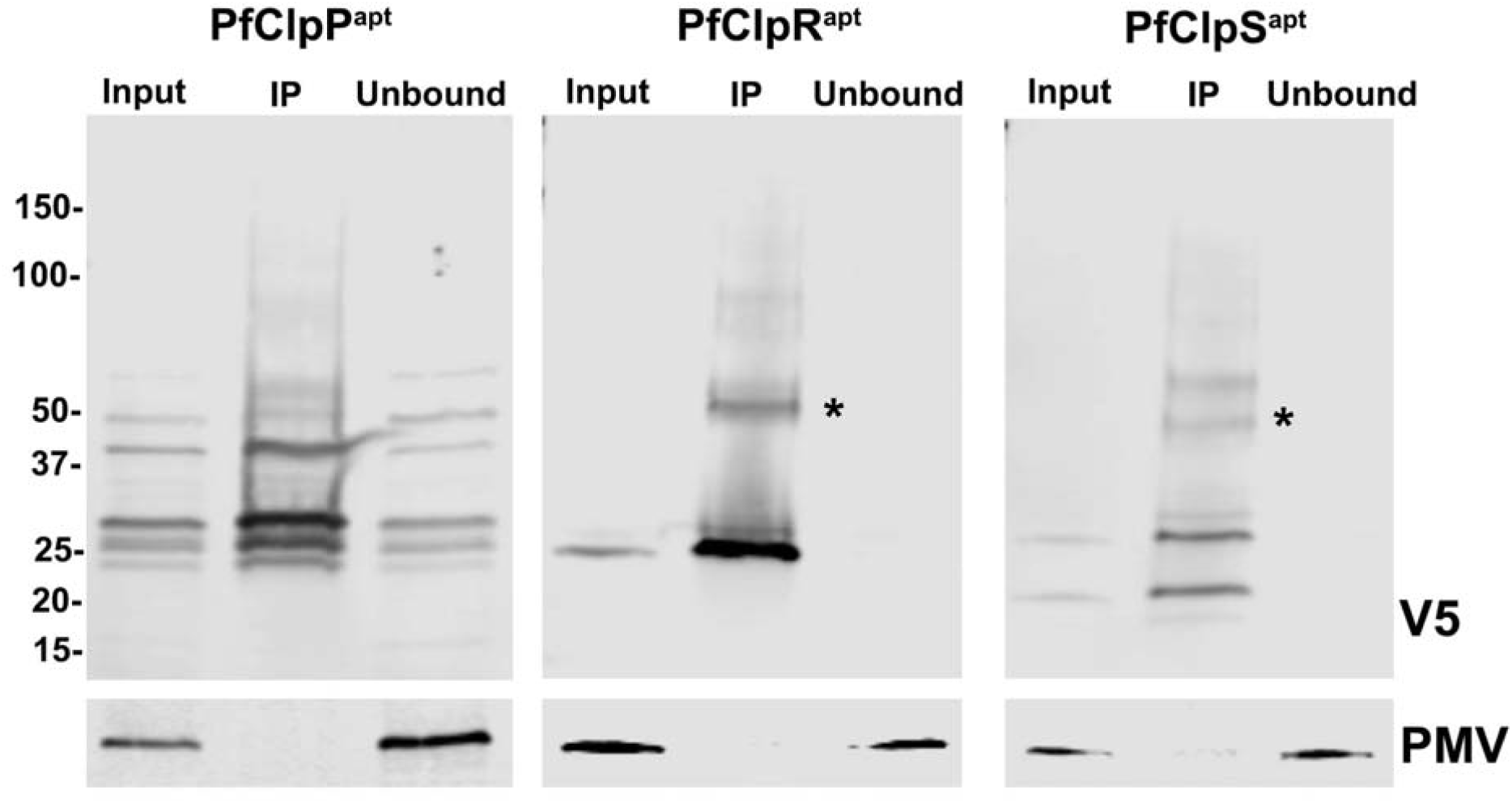
Immunoprecipitation of PfClpP^apt^, PfClpR^apt^ and PfClpS^apt^ for proteomic analysis. 10^9^ parasites were isolated and 10% of the volume was saved as an input sample. Immunoprecipitation of the tagged proteins from respective parasite lysates were performed using anti-V5 antibody-conjugated beads. Input, IP and unbound samples were separated on SDS-PAGE and blotted with anti-V5 antibodies and anti-PMV (Plasmpesin V, loading control). The heavy chain of the antibody that eluted with some of the IP samples, despite cross linking is shown (*).

## Notes

#### Summary of Updates

Updated title and some corrections were made.

